# Positions of pivot points in quadrupedal locomotion: limbs and trunk global control in four different dog breeds

**DOI:** 10.1101/2022.12.09.519601

**Authors:** Emanuel Andrada, Gregor Hildebrandt, Hartmut Witte, Martin S. Fischer

## Abstract

Dogs (*Canis familiaris*) prefer the walk at lower speeds and the more economical trot at speeds ranging from 0.5 Fr up to 3 Fr. Important works were carried out to understand these gaits at the levels of center of mass, joint mechanics, and muscular control. However, less is known about the global control goals for limbs and overall locomotion, and of whether these global control goals are gait or breed specific. For walk and trot, we analyzed dog global dynamics based on motion capture and single leg kinetic data recorded from treadmill locomotion of French Bulldog (N = 4), Whippet (N = 5), Malinois (N = 4) and Beagle (N = 5). Dogs displayed two virtual pivot points (VPP) during walk and trot each. One resembles control of both thoracic (fore) limbs and is roughly located above and caudally to the scapular pivot, while the second is located roughly above and cranially to the hip and mirrors the control of the pelvic (hind-) limbs. The positions of VPPs and the patterns of the legs‘ axial and tangential functions were gait and breed related. However, breed related changes were mainly exposed by the French Bulldog. The position of VPPs relative to the proximal pivots explains the propulsive and breaking forces observed in quadrupedal locomotion and may help to reduce limb work. In combination with former work, from the present study the VPP template emerges as the expression of a simple and general global control rule for both bipeds and quadrupeds.

## Introduction

Dogs prefer the walk at lower speeds and the (more economical) trot at speeds ranging from 0.5 Fr up to 3 Fr (Bryce and Williams, 2017; Jayes and Alexander, 1978). Fr is a dimensionless measure of speed known as the Froude number (*Fr* = *v*_*t*_^2^/*gl*), where *v*_*t*_ is the locomotion speed, *g* is gravitational acceleration, *l* is the effective leg length (segment between most proximal leg pivot and the ground contact point, see Fig. 1). Walking dogs alternate between a short 2-legged and long 3-legged support of the body, trotting dogs use diagonal pairs of limbs. Different leg coordination is mirrored in the mechanics of the center of mass, i.e. vaulting mechanics (Cavagna et al., 1977) at walk vs. bouncing mechanics (Blickhan, 1989) at trot. Yet, gait related differences are more diffuse when looking at the level of joint dynamics or muscle activations (Andrada et al., 2017; Stark et al., 2021; Tokuriki, 1973a; Tokuriki, 1973b).

**Figure 1.**
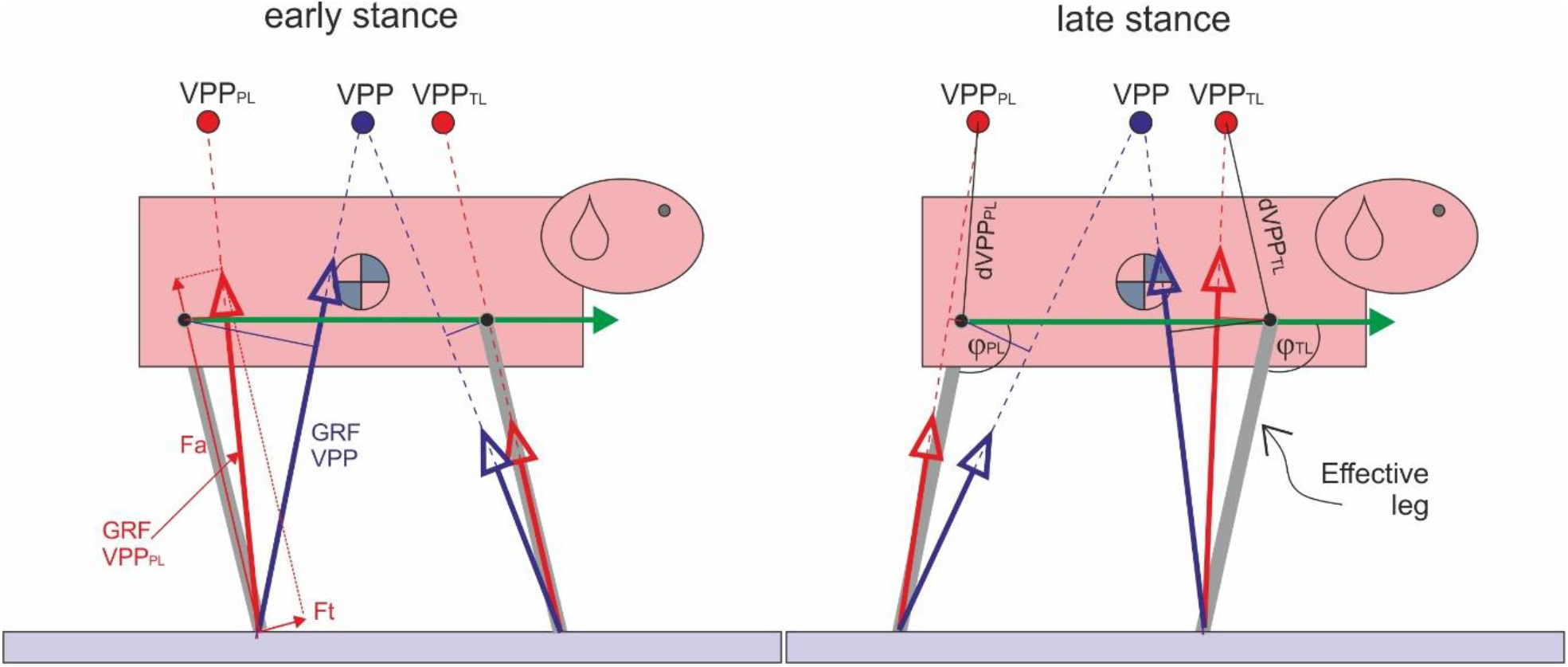
Influence of number of VPPs on pelvic and thoracic effective leg mechanics. Controlling locomotion in quadrupeds with one VPP above the CoM as observed in bipeds would induce large joint torque and work in the proximal pivots due to the larger lever arms (thin blue solid lines). Moreover, pelvic and thoracic limbs cannot be controlled independently. Two proximal joint related VPPs (VPP_PL_ and VPP_TL_) may solve these problems. Green arrow represents trunk vector, φ_TL_ and φ_PL_: angles between trunk vector and effective legs. Fa: axial force, Ft: tangential force, PL: pelvic limb, TL: thoracic limb, dVPP_TL_: distance VPP_PL_-hip joint), dVPP_TL_: distance VPP_PL_-scapular pivot (center of scapular rotation in a sagittal plane).

Important work has been done to understand dog locomotion at the muscular (Carrier et al., 2006; Carrier et al., 2008; Deban et al., 2012; Goslow et al., 1981; Tokuriki, 1973a; Tokuriki, 1973b; Tokuriki, 1974) and at joint levels in healthy and sick dogs (Andrada et al., 2017; Burton et al., 2008; Burton et al., 2011; Carrier et al., 1998; Colborne et al., 2011; Colborne et al., 2005; Colborne et al., 2006; Dogan et al., 1991; Gregersen et al., 1998; Headrick, 2012; Headrick et al., 2014; Nielsen et al., 2003; Ragetly et al., 2010). Still, we know little about the global control goals of limbs for periodic/stable locomotion. In addition, it is unknown whether these global control goals are specific for gait or breed. If global limb control among different breeds is conservative, then differences in limb kinematics, e.g. (Fischer et al., 2018; Fischer and Lilje, 2011) might reflect adaptations to body-, limb proportions and posture.

At the global limb level, the thoracic limbs (fore limbs) have a primordial weight-bearing function and contribute less to propulsion than do the pelvic limbs (hindlimbs) (Bertram et al., 2000; Budsberg et al., 1987; Lee et al., 1999). Based on the absence of (or extremely low) activity of the main protractor and retractor muscles of the humerus during the stance phase, Carrier and colleagues hypothesized that the thoracic limbs mainly work axially (i.e., as struts at the shoulder joints) (Carrier et al., 2008). This description agrees with the spring-loaded inverted pendulum (SLIP) model. Despite its simplicity, template models like the SLIP model allow to extract key features of quadrupedal locomotion. With such a simple representation, McMahon could explain why galloping is a faster quadrupedal gait than trotting (McMahon, 1985). With a model composed of a rigid torso and prismatic, massless, spring-like effective legs, other authors analyzed the energetics of trotting, bounding and galloping (Nanua, 1992; Nanua and Waldron, 1995). Later, Poulakakis and colleagues used a similar model to analyze locomotion stability in the sagittal plane (Poulakakis et al., 2003; Poulakakis et al., 2006). Other extensions of those simple models included an articulated torso e.g., (Cao and Poulakakis, 2013; Deng et al., 2012). Recently, Söhnel et al., presented global leg functions of jumping dogs. They could separate beginner from skilled agility dogs based on SLIP-related experimental parameters (Söhnel et al., 2020).

In the last years, there is an increasing consent among scientists, that the global behavior of the leg, represented by the effective leg, can be separated into two main time-related functions: an axial and a tangential or rotational leg function e.g., (Andrada et al., 2014; Maus et al., 2010; Shen and Seipel, 2012). The axial leg function combines the axial force (*F*_*a*_) with the length change of the instantaneous effective leg (*F*_*a*_ vs. Δ*l*) relative to the leg length at TD (*l*_0_). Fa is the component of the ground reaction force (GRF) along the effective leg (axial). The axial leg function provides mainly a weight bearing function, generates vertical body oscillations and is usually represented by leg stiffness, or more exactly leg impedance.

The tangential leg function can be displayed by combining the proximal torque with the joint angle (*M* vs φ). The proximal joint torque *M* is obtained by multiplying the force perpendicular to the effective leg (*F*_*t*_, therefore tangential function) by the instantaneous effective leg length *l*. The tangential leg function represents the control strategy used to retract the effective leg and to balance the trunk. The vectorial sum of *F*_*a*_ and *F*_*t*_ yields the vector of GRF which we measure via force plates (see Fig. 1).

To generate periodic locomotion, both axial and tangential leg functions must be combined in a way that leg retraction matches the oscillation time along the leg. Experimental evidence shows that in bipeds such as humans and birds (Andrada et al., 2014; Maus et al., 2010) and also facultative bipeds such as Japanese macaques (Blickhan et al., 2018) axial and tangential leg functions are related to each other via a “Virtual Pivot Point” (VPP). The VPP is a body-fixed point, mostly located above the center of mass, at which the GRF vector points during stance. The term “virtual” indicates that the body rotates around a pivot point that is not an anatomical joint. In bipeds, VPP as the target of control simplifies the balance of the trunk by changing an inverted pendulum control problem into a suspended one. Moreover, simulation studies based on this bio-inspired strategy showed that VPP control can balance not only the trunk but also generate stable bipedal gaits (Andrada et al., 2014; Drama and Badri-Spröwitz, 2020; Maus et al., 2010). So far, the existence of a VPP in quadrupedal locomotion has not been demonstrated experimentally. Maus and colleagues searched for one VPP for all four legs on a dog at trot, but results were unclear (Maus et al., 2010). Japanese macaques displayed in experiments after just few months of bipedal training a body-fixed VPP (Blickhan et al., 2018). A simple explanation would be that they already had been using such a control scheme during quadrupedal locomotion. However, the use of one body fixed VPP in quadrupeds would be disadvantageous at least in two ways: first, pelvic and thoracic limbs would not be independent from each other and second, joint torque and work in the most proximal joints would be quite large. Interestingly, these two problems may be solved if each pair of limbs has a proximal joint related VPP (see Fig. 1). With this background, we phrased the following *hypotheses*:

1. After accounting for mass and length measures, control principles of locomotion at the global level are similar among different dog breeds.
2. If (1) is falsified, then breed related differences in the position of the VPPs and or in the leg axial and tangential leg functions may inform about limb global control adaptations related to body proportions, posture, and behavior.

## Materials & Methods

### Animals

The present work uses a subset of the data collected for (Andrada et al., 2017) and for (Fischer et al., 2018). Animals details were published in those works and will be only briefly summarized here: we collected data from five adult male Beagles belonging to a research colony based at the Small Animal Hospital of the University of Veterinary Medicine, Hannover, Germany, four adult Malinois (3 males/1 female) kept as police dogs by the Saxonian police force, four adult female French Bulldogs from private dog owners, and five adult Whippets (2 males/3 females) from a private dog owner. Table 1 describes the available material.

**Table 1:** Dogs and number of steps analyzed for this study.

| Breed | Individual | W [kg] | HW [cm] | Steps walk |  | Steps trot |  |
| --- | --- | --- | --- | --- | --- | --- | --- |
|  |  |  |  | PL | TL | PL | TL |
| Beagle | Erwin | 14.9 | 35 | 10 | 10 | 13 | 12 |
|  | Simon | 13.8 | 33 | 15 | 16 | 9 | 8 |
|  | Malte | 14.8 | 34 | 26 | 26 | 32 | 31 |
|  | Louis | 16.2 | 38 | 21 | 14 | 21 | 21 |
|  | Spencer | 19.8 | 42 | 12 | 11 | 29 | 13 |
| Malinois | Zora | 28.5 | 64 | 9 | 12 | 9 | 5 |
|  | Pike | 28.5 | 62 | 9 | 10 | 5 | 8 |
|  | Hunter | 18.6 | 59 | 10 | 10 | 10 | 7 |
|  | Rocky | 22.4 | 58 | 7 | 10 | 10 | 8 |
| French Bulldog | Queny | 11 | 31 | 15 | 17 | 20 | 15 |
|  | MJ | 9,5 | 30 | 17 | 16 | 18 | 7 |
|  | Chacha | 10 | 31 | 19 | 19 | 18 | 8 |
|  | Juno | 13 | 32 | 20 | 16 | 19 | 19 |
| Whippet | Lilly | 12 | 49 | 18 | 19 | 19 | 5 |
|  | Kenja | 10 | 46 | 4 | 3 | 21 | 20 |
|  | Afrika | 9 | 47 | 16 | 15 | 13 | 41 |
|  | Merlin | 16.5 | 51 | 18 | 17 | 10 | 6 |
|  | Moody | 13.3 | 50 | 17 | 20 | 12 | 6 |

W: weight, HW: height at the withers, PL: pelvic limb, TL: thoracic limb

### Marker setup, motion analysis and kinetics

The marker setup encompassed 19 markers on the left pelvic limb and 21 markers on the left pelvic limb. For the propose of this work we used only the most proximal and distal leg markers to describe both the pelvic and the thoracic effective legs and the trunk. For the pelvic limb the effective leg was computed as the direct connection between the marker placed on the dorsal aspect of the of the third metatarsal bones and the marker located at the greater trochanter of the femur. Thoracic limb effective leg was computed as the distance between the dorsal aspect of the third phalanx and the scapular pivot. Based on our previous works we assumed that the scapular pivot is located at 2/3 of the distance between the markers representing the most dorsal and ventral points of the scapula (Andrada et al., 2017; Fischer, 1999). The trunk was defined as a vector from hip joint (greater trochanter of the femur) to the scapular pivot.

3D kinematic data was collected using 6 infrared Vicon® cameras (Oxford Metrics, Oxford, UK) and an instrumented quad-band treadmill (model 4060-08, Bertec Corporation) available at the locomotion lab of the Small Animal Hospital of the University of Veterinary Medicine Hannover, Germany. Kinematic data was collected at 100 Hz. Data collection started as soon as the dogs were walking or trotting smoothly and comfortably. Data was recorded for a maximum of 45 s. For computation, series of at least 5 cycles (strides) in which the dog moved steadily and without overstepping onto the other bands (force plates) were used. When trotting, dogs were kept on one side of the treadmill (usually the left side) to facilitate handling. Computed number of steps can be found in Table 1. The lab coordinate system was set as follows: +x pointed left, +y pointed opposite to the direction of motion and +z pointed upwards. The 3D coordinates of marker trajectories were smoothed by a Butterworth four order low-pass filter with a cut-off frequency of 6 Hz.

In this work we used the sagittal projection of both kinematic and GRF. To describe the axial leg function, we combined the instant changes of effective leg length 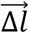 and axial force 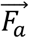 during stance.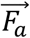 was computed by projecting the vector of the GRF into the vector defining the effective leg 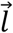 (see Fig. 1). The tangential function combines joint torque *M* and effective leg angle φ relative to the trunk vector. The trunk vector was defined as a vector between hip joint and scapular pivot. The angle φ between trunk vector and effective leg was computed using the dot product between two vectors (see Fig. 1). Proximal joint torque was computed as 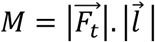 were the tangential force 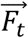 was previously obtained by computing the component of the vector of the GRF perpendicular to 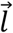 (see Fig. 1). Axial work was computed as the area inside the loop in the graph 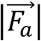 vs 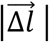, while tangential work as the area below the curve *M* vs φ. VPP and paw positions were computed relative to the proximal pivots (hip and scapular pivot) adapting the methods proposed for the CoM in (Andrada et al., 2014). Note that for each limb pair, we described the relative position of the distal point of the effective legs and the direction of the GRF during the stance from a moving frame. Therefore, although e.g., the scapular pivot translates relative to the trunk during stance in global coordinates, in the relative coordinates we used it is a fixed point. In plots, the positions of the proximal pivots were frozen at their values at TD. For the sake of comparison, we transformed force, length, and torque into a nondimensional form. For the force we used 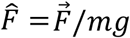 for the length 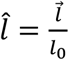, being *l*_0_ the effective leg length at TD, for the change of leg length 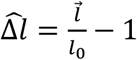, and for the torque 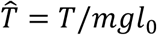. Global dynamics were computed using a custom written script in Matlab 2017 (The MathWorks Inc., Natick, MA, USA).

### Statistical analysis

To infer the influence of gait and breed on the VPP position, maximal axial force *F*_a-max_, maximal joint Torque *M*_max_, and leg angle relative to the trunk at TD (φ_0_), repeated measures ANOVA with Gait (walk vs. trot) as within subjects and breed as between subjects were performed. Afterwards, Post hoc tests with Bonferroni correction were performed. Statistical analysis was performed in IBM® SPSS® Statistics 26.

## Results

All breeds displayed a proximal joint related VPP point above both the hip and the scapular pivots (center of scapular rotation in the sagittal plane) during both walk and trot. In addition, our results show that leg function is rather similar among different dog breeds, but for French Bulldogs and Whippets some deviations were found.

### Pelvic limb, axial leg function

At walk, the axial leg function diverges from the spring-like leg behavior during stance. Malinois and Beagles displayed similar leg functions. Both exhibited effective leg’s lengthening of about 4% with respect to the length measured at TD. A picture closer to a spring-like leg behavior was observed for Whippets, although they also showed only small leg lengthening (Figure 2). Contrarily, in French Bulldogs leg length was shortened during stance to about the same amount.

**Figure 2.**
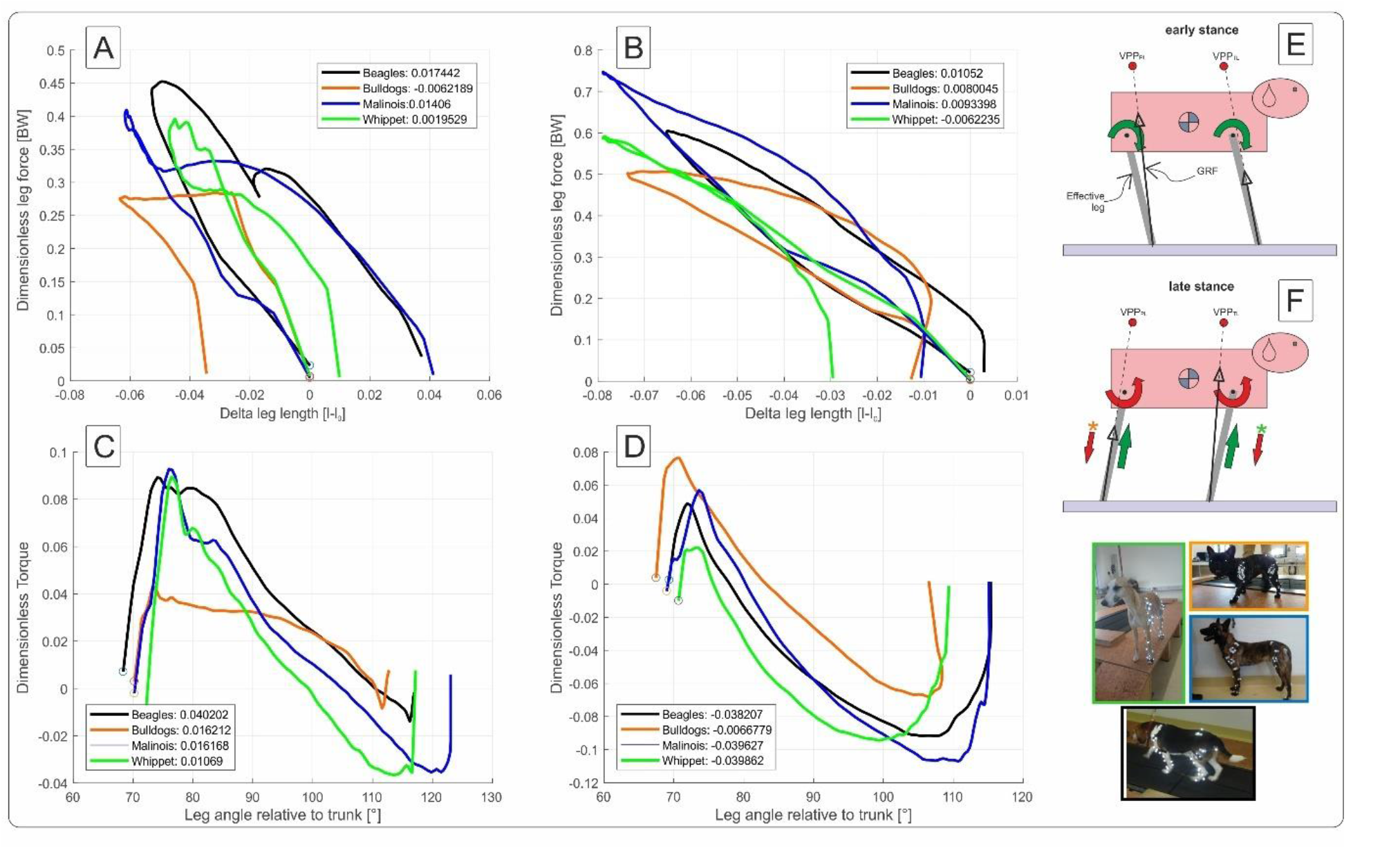
Axial and tangential leg functions at walk in Beagle (black), Whippet (green), French Bulldog (orange) and Malinois (blue). A, C: pelvic limb, B, D: thoracic limb. A, B: axial function and C, D: tangential function of the effective leg. Curves represent mean values. Positive values represent retractor, while negative protractor torques. Legends show net axial/tangential work. E, F: template representation at early and late stance based on the curves in A, B, C, D. Position of the VPPs and leg orientations are rough approximations. For more accurate positions see Fig. 4 and Tab. 4. In E and F, curved arrows represent torque direction. Linear arrows indicate leg extension/shortening. Green arrows indicate energy generation (motions and force/torque directions coincide), red energy absorption (motions and force/torque directions are opposite). Note that in F arrows with (*) display generalized leg functions in French Bulldog and Whippet that differed from the two other breeds.

The leg lengthening was gait related (p < 0.001) but not breed related. At trot, all four breeds exhibited leg lengthening during late stance phase (Fig. 3 and Tab. 2). Again, Beagles and Malinois displayed similar leg functions. They showed the highest projected leg force (F_ax_ > 0.7) and leg lengthening between 6% and 7%, and thus the highest positive axial work (see Fig 3A). Whippets displayed lower peak axial forces and less leg lengthening than Malinois and Beagles. French Bulldogs displayed the lowest peak axial force (about 0.5 BW) and a more spring-like leg behavior. Therefore, they exerted the lowest positive net axial work.

**Figure 3.**
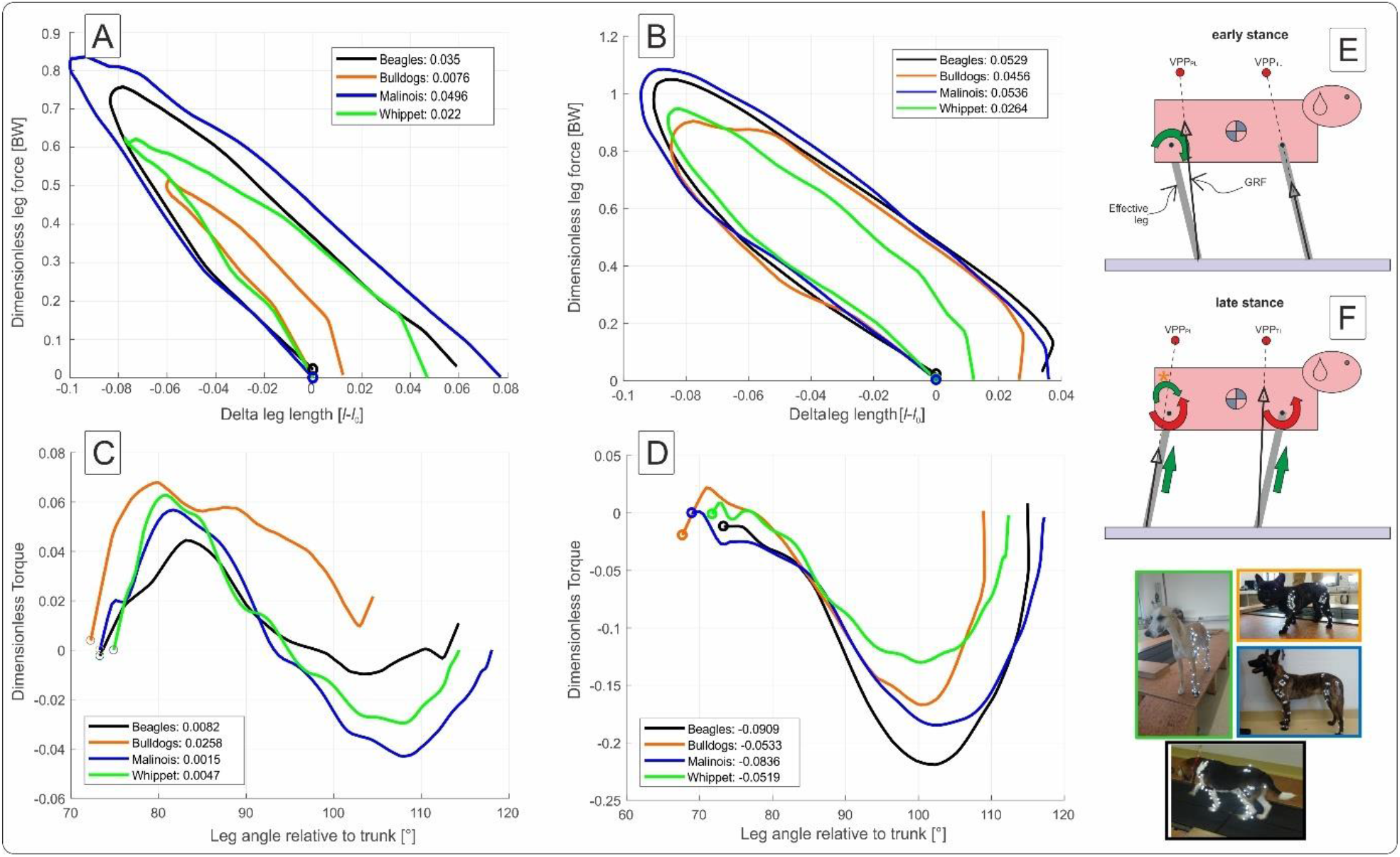
Axial and tangential leg functions at trot in Beagle (black), Whippet (green), French Bulldog (orange) and Malinois (blue). A, C: pelvic limb, B, D: thoracic limb. A, B: axial function and C, D: tangential function of the effective leg. Curves represent mean values. Positive values represent retractor while negative protractor torques. Legends show net axial/tangential work. E, F: template representation at early and late stance based on the curves A, B, C, D. Position of the VPPs and leg orientations are roughly approximations and may vary between breeds. For accurate data see Fig. 4 and Tab. 4. In E and F, curved arrows represent torque direction. Linear arrows indicate leg extension/shortening. Green arrows indicate energy generation (motions and force/torque directions coincide), red energy absorption (motions and force/torque directions are opposite). Note that in F the curved arrow with (*) display tangential leg function in the French Bulldog that differed from the mainstream.

**Table 2:** Pelvic limb: components of the axial (Fa_PL_ vs. Δl_PL_) and of the tangential leg function (*M*PL vs φ_PL_)

| % of stance time | 1 | 25 | 50 | 75 | 99 |
| --- | --- | --- | --- | --- | --- |
| **Beagle** |  |  |  |  |  |
| $\Delta l_{PL}$ (w) | $-0.01 \pm 0$ | $-0.05 \pm 0.04$ | $-0.02 \pm 0.04$ | $-0.02 \pm 0.03$ | $0.03 \pm 0.03$ |
| $\Delta l_{PL}$ (t) | $-0.003 \pm 0$ | $-0.071 \pm 0.01$ | $-0.074 \pm 0.02$ | $-0.010 \pm 0.013$ | $0.057 \pm 0.01$ |
| $F_{aPL}$ (w) | $0.06 \pm 0$ | $0.45 \pm 0.03$ | $0.29 \pm 0.03$ | $0.32 \pm 0.03$ | $0.06 \pm 0.02$ |
| $F_{aPL}$ (t) | $0.042 \pm 0.001$ | $0.552 \pm 0.009$ | $0.747 \pm 0.016$ | $0.426 \pm 0.013$ | $0.044 \pm 0.009$ |
| $\varphi_{PL}$ (w) [°] | $68.9 \pm 1.4$ | $80.1 \pm 2.7$ | $92.1 \pm 3.1$ | $106.6 \pm 3.5$ | $116.7 \pm 3.5$ |
| $\varphi_{PL}$ (t) [°] | $73.7 \pm 1.5$ | $85.3 \pm 1.7$ | $97.6 \pm 2.5$ | $106.8 \pm 4.1$ | $114.0 \pm 4.2$ |
| $M_{PL}$ (w) | $0.020 \pm 0.005$ | $0.085 \pm 0.037$ | $0.043 \pm 0.021$ | $0.009 \pm 0.015$ | $-0.007 \pm 0.009$ |
| $M_{PL}$ (t) | $0.001 \pm 0.006$ | $0.041 \pm 0.014$ | $0.001 \pm 0.022$ | $-0.005 \pm 0.013$ | $0.009 \pm 0.008$ |
| **French Bulldog** |  |  |  |  |  |
| $\Delta l_{PL}$ (w) | $-0.003 \pm 0$ | $-0.028 \pm 0.02$ | $-0.053 \pm 0.026$ | $-0.062 \pm 0.033$ | $-0.036 \pm 0.04$ |
| $\Delta l_{PL}$ (t) | $-0.002 \pm 0$ | $-0.047 \pm 0.01$ | $-0.058 \pm 0.014$ | $-0.024 \pm 0.01$ | $0.011 \pm 0.012$ |
| $F_{aPL}$ (w) | $0.04 \pm 0$ | $0.28 \pm 0.02$ | $0.28 \pm 0.03$ | $0.27 \pm 0.03$ | $0.05 \pm 0.033$ |
| $F_{aPL}$ (t) | $0.03 \pm 0$ | $0.40 \pm 0.01$ | $0.51 \pm 0.01$ | $0.36 \pm 0.01$ | $0.03 \pm 0.014$ |
| $\varphi_{PL}$ (w) [°] | $70.5 \pm 3.8$ | $76.5 \pm 3.6$ | $87.3 \pm 2.6$ | $100.8 \pm 1.74$ | $112.5 \pm 3.6$ |
| $\varphi_{PL}$ (t) [°] | $72.6 \pm 3.2$ | $81.0 \pm 3$ | $89.9 \pm 2.9$ | $97.9 \pm 2.6$ | $104.3 \pm 2.9$ |
| $M_{PL}$ (w) | $0.010 \pm 0.027$ | $0.038 \pm 0.106$ | $0.033 \pm 0.074$ | $0.022 \pm 0.034$ | $0.005 \pm 0.008$ |
| $M_{PL}$ (t) | $-0.012 \pm 0.012$ | $-0.075 \pm 0.014$ | $-0.218 \pm 0.01$ | $-0.144 \pm 0.01$ | $-0.002 \pm 0.006$ |
| **Malinois** |  |  |  |  |  |
| $\Delta l_{PL}$ (w) | $-0.006 \pm 0$ | $-0.061 \pm 0.025$ | $-0.052 \pm 0.02$ | $-0.028 \pm 0.031$ | $0.040 \pm 0.028$ |
| $\Delta l_{PL}$ (t) | $-0.004 \pm 0$ | $-0.085 \pm 0.013$ | $-0.09 \pm 0.02$ | $-0.011 \pm 0.02$ | $0.075 \pm 0.019$ |
| $F_{aPL}$ (w) | $0.05 \pm 0$ | $0.41 \pm 0.021$ | $0.33 \pm 0.022$ | $0.33 \pm 0.06$ | $0.04 \pm 0.02$ |
| $F_{aPL}$ (t) | $0.03 \pm 0$ | $0.65 \pm 0.01$ | $0.83 \pm 0.02$ | $0.51 \pm 0.022$ | $0.02 \pm 0.01$ |
| $\varphi_{PL}$ (w) [°] | $70.9 \pm 3.8$ | $83.3 \pm 4.7$ | $97.0 \pm 5.2$ | $113.0 \pm 5.1$ | $123.0 \pm 7.5$ |
| $\varphi_{PL}$ (t) [°] | $73.8 \pm 3.2$ | $86.4 \pm 3.5$ | $100.1 \pm 4.1$ | $111.1 \pm 5.2$ | $117.9 \pm 5.7$ |
| $M_{PL}$ (w) | $0.012 \pm 0.015$ | $0.063 \pm 0.028$ | $0.016 \pm 0.023$ | $-0.021 \pm 0.024$ | $-0.008 \pm 0.013$ |
| $M_{PL}$ (t) | $0.007 \pm 0.012$ | $0.046 \pm 0.014$ | $-0.025 \pm 0.01$ | $-0.036 \pm 0.01$ | $-0.002 \pm 0.006$ |
| **Whippet** |  |  |  |  |  |
| $\Delta l_{PL}$ (w) | $-0.004 \pm 0$ | $-0.045 \pm 0.02$ | $-0.044 \pm 0.026$ | $-0.030 \pm 0.022$ | $0.009 \pm 0.033$ |
| $\Delta l_{PL}$ (t) | $-0.003 \pm 0$ | $-0.063 \pm 0.011$ | $-0.073 \pm 0.021$ | $-0.021 \pm 0.026$ | $0.045 \pm 0.025$ |
| $F_{aPL}$ (w) | $0.06 \pm 0$ | $0.39 \pm 0.02$ | $0.32 \pm 0.021$ | $0.29 \pm 0.01$ | $0.05 \pm 0.03$ |
| $F_{aPL}$ (t) | $0.03 \pm 0$ | $0.55 \pm 0.01$ | $0.62 \pm 0.05$ | $0.45 \pm 0.02$ | $0.03 \pm 0.02$ |
| $\varphi_{PL}$ (w) [°] | $72.6 \pm 3.1$ | $82.8 \pm 3.2$ | $93.5 \pm 3.3$ | $106.1 \pm 5$ | $117.0 \pm 5.6$ |
| $\varphi_{PL}$ (t) [°] | $75.3 \pm 2$ | $85.6 \pm 3$ | $97.6 \pm 4.2$ | $108.0 \pm 5$ | $114.2 \pm 4.4$ |
| $M_{PL}$ (w) | $0.005 \pm 0.016$ | $0.055 \pm 0.012$ | $0.014 \pm 0.014$ | $-0.023 \pm 0.023$ | $-0.008 \pm 0.005$ |
| $M_{PL}$ (t) | $0.007 \pm 0.007$ | $0.048 \pm 0.022$ | $-0.010 \pm 0.035$ | $-0.030 \pm 0.030$ | $-0.001 \pm 0.009$ |

Mean values ± SD for the axial force ( $F_{aPL} = Fa/mg$ ), the leg length change ( $\Delta l_{PL} = l/l_0 - 1$ ), the angle between leg and trunk vector ( $\varphi_{PL}$ ), and the hip torque ( $M_{PL}$ ) at walk (w) and at trot (t). Note that $F_{aPL}$ , $\Delta l_{PL}$ and $M_{PL}$ are dimensionless. $l$ is leg length and $l_0$ leg length at TD, $m$ = mass and $g$ = gravity.

### Pelvic limb, tangential leg function

At walk the angle of attack φ_0_ is at about 70° in all four breeds, and the lift off angle φ between 112° (French Bulldog) and 123° (Malinois), see Table 2. During most of the stance phase the hip muscles actively retract the pelvic limb (positive torque and leg retraction, Fig. 2C).

Maximal positive peak torque was similar in time and value (T∼ 0.09) for Whippets, Malinois and Beagles, but about the half of that value for French Bulldogs. The mean torque profile was basically biphasic. However, only Malinois and Whippets displayed a markedly negative torque (leading to protraction) in the late stance phase.

During trot, effective legs touched the ground a steeper than at walk (p < 0.001). Effective legs were less retracted during stance at trot compared to walk (p < 0.05). Mean torque profiles were biphasic for all analyzed breeds except for French Bulldogs, which showed a half sinus profile (Fig. 3C). Maximal Torques were gait related (p < 0.05). French Bulldogs also displayed the largest peak positive torque. For the other breeds, the maximal positive (retractor) torques were lower than those exhibited during walk.

### Pelvic limbs, VPP

The position of the pelvic limbs’ VPP (VPP_PL_) is gait related. The VPP_PL_ point was significantly (p < 0.001) higher placed at walk (about 0.5 and 0.9 of leg length at TD) compared to its position at trot (close to 0.2 of leg length at TD). However, the distance between VPP_PL_ and hip joint did not significantly vary between breeds nor did the linear combination of gait effects * breed effects (see Tab. 4).

### Thoracic limb, axial leg function

At walk the axial leg function displayed a decent spring-like leg behavior during stance. In Whippet only the last 10% of stance diverged from a worthy spring-like leg and finished with a leg shortening of about 3% of *l*_0_ and negative axial work. The other breeds, even when showing very small leg lengthening or shortening, produced positive net axial work (see Fig. 2B and Tab. 3). At trot, all breeds displayed similar axial leg functions, and exhibited leg enlargement (between 1.1% for Whippets, 2.7% for French Bulldogs and about 3.5% for Beagles and Malinois). Consequently, all breeds generated positive net axial work in the thoracic limb (see Fig. 3B and Tab. 3).

**Table 3:** Thoracic limb: components of the axial (Fa_TL_ vs. Δl_TL_) and of the tangential leg function (*M*TL vs φ_TL_)

| % stance phase | 1 | 25 | 50 | 75 | 99 |
| --- | --- | --- | --- | --- | --- |
| **Beagle** |  |  |  |  |  |
| $\Delta l_{TL}$ (w) | $-0.004 \pm 0$ | $-0.064 \pm 0.01$ | $-0.055 \pm 0.026$ | $-0.034 \pm 0.035$ | $0.003 \pm 0.04$ |
| $\Delta l_{TL}$ (t) | $-0.004 \pm 0$ | $-0.077 \pm 0.01$ | $-0.079 \pm 0.014$ | $-0.005 \pm 0.015$ | $0.035 \pm 0.02$ |
| $F_{aTL}$ (w) | $0.06 \pm 0$ | $0.58 \pm 0.01$ | $0.57 \pm 0.02$ | $0.43 \pm 0.03$ | $0.06 \pm 0.05$ |
| $F_{aTL}$ (t) | $0.05 \pm 0$ | $0.67 \pm 0.01$ | $1.04 \pm 0.01$ | $0.53 \pm 0.01$ | $0.05 \pm 0.02$ |
| $\varphi_{TL}$ (w) [°] | $69.6 \pm 3.4$ | $80.3 \pm 1.7$ | $96.7 \pm 1.8$ | $108.5 \pm 2$ | $115.4 \pm 2.9$ |
| $\varphi_{TL}$ (t) [°] | $73.8 \pm 5.3$ | $86.6 \pm 4.4$ | $100.8 \pm$ | $111.5 \pm 3.3$ | $115 \pm 3$ |
| $M_{TL}$ (w) | $0.01 \pm 0.01$ | $-0.02 \pm 0.03$ | $-0.08 \pm 0.04$ | $-0.09 \pm 0.03$ | $-0.01 \pm 0.03$ |
| $M_{TL}$ (t) | $-0.012 \pm 0.01$ | $-0.075 \pm 0.003$ | $-0.218 \pm 0.05$ | $-0.144 \pm 0.028$ | $-0.002 \pm 0.01$ |
| **French Bulldog** |  |  |  |  |  |
| $\Delta l_{TL}$ (w) | $-0.004 \pm 0$ | $-0.063 \pm 0.014$ | $-0.066 \pm 0.029$ | $-0.032 \pm 0.036$ | $-0.012 \pm 0.035$ |
| $\Delta l_{TL}$ (t) | $-0.004 \pm 0$ | $-0.076 \pm 0.01$ | $-0.072 \pm 0.02$ | $-0.011 \pm 0.024$ | $0.027 \pm 0.024$ |
| $F_{aTL}$ (w) | $0.05 \pm 0$ | $0.45 \pm 0.01$ | $0.51 \pm 0.08$ | $0.42 \pm 0.03$ | $0.05 \pm 0.03$ |
| $F_{aTL}$ (t) | $0.04 \pm 0$ | $0.63 \pm 0.01$ | $0.89 \pm 0.02$ | $0.55 \pm 0.02$ | $0.03 \pm 0.02$ |
| $\varphi_{TL}$ (w) [°] | $67.8 \pm 5$ | $77.5 \pm 4.4$ | $92.4 \pm 3.8$ | $104.6 \pm 3.5$ | $106.9 \pm 5.1$ |
| $\varphi_{TL}$ (t) [°] | $68.2 \pm 5$ | $82.0 \pm 5.4$ | $95.5 \pm 4.7$ | $105.3 \pm 4.2$ | $109.0 \pm 2.8$ |
| $M_{TL}$ (w) | $0.024 \pm 0.007$ | $0.032 \pm 0.032$ | $-0.035 \pm 0.036$ | $-0.067 \pm 0.032$ | $-0.011 \pm 0.005$ |
| $M_{TL}$ (t) | $-0.01 \pm 0.01$ | $-0.03 \pm 0.04$ | $-0.15 \pm 0.09$ | $-0.13 \pm 0.06$ | $-0.01 \pm 0.01$ |
| **Malinois** |  |  |  |  |  |
| $\Delta l_{TL}$ (w) | $-0.004 \pm 0$ | $-0.074 \pm 0.01$ | $-0.073 \pm 0.004$ | $-0.043 \pm 0.01$ | $-0.010 \pm 0.023$ |
| $\Delta l_{TL}$ (t) | $-0.004 \pm 0$ | $-0.083 \pm 0.005$ | $-0.084 \pm 0.006$ | $-0.019 \pm 0.006$ | $0.036 \pm 0.009$ |
| $F_{aTL}$ (w) | $0.05 \pm 0$ | $0.70 \pm 0.01$ | $0.70 \pm 0$ | $0.55 \pm 0.01$ | $0.05 \pm 0.02$ |
| $F_{aTL}$ (t) | $0.04 \pm 0$ | $0.72 \pm 0.01$ | $1.08 \pm 0.01$ | $0.65 \pm 0.01$ | $0.03 \pm 0.01$ |
| $\varphi_{TL}$ (w) [°] | $69.3 \pm 1.9$ | $81.2 \pm 3.2$ | $97.0 \pm 4.1$ | $108.9 \pm 2.5$ | $115.1 \pm 2.3$ |
| $\varphi_{TL}$ (t) [°] | $69.5 \pm 4.4$ | $84.6 \pm 3.4$ | $100.7 \pm 3.1$ | $111.7 \pm 3.6$ | $117.2 \pm 2.8$ |
| $M_{TL}$ (w) | $0.005 \pm 0.006$ | $-0.007 \pm 0.020$ | $-0.083 \pm 0.025$ | $-0.106 \pm 0.046$ | $0.005 \pm 0.014$ |
| $M_{TL}$ (t) | $0.001 \pm 0.016$ | $-0.051 \pm 0.038$ | $-0.182 \pm 0.049$ | $-0.135 \pm 0.050$ | $-0.006 \pm 0.005$ |
| **Whippet** |  |  |  |  |  |
| $\Delta l_{TL}$ (w) | $-0.004 \pm 0$ | $-0.070 \pm 0.014$ | $-0.076 \pm 0.017$ | $-0.051 \pm 0.017$ | $-0.030 \pm 0.016$ |
| $\Delta l_{TL}$ (t) | $-0.004 \pm 0$ | $-0.073 \pm 0.006$ | $-0.080 \pm 0.007$ | $-0.032 \pm 0.006$ | $0.011 \pm 0.003$ |
| $F_{aTL}$ (w) | $0.05 \pm 0$ | $0.54 \pm 0.01$ | $0.58 \pm 0.02$ | $0.43 \pm 0.02$ | $0.06 \pm 0.02$ |
| $F_{aTL}$ (t) | $0.04 \pm 0$ | $0.67 \pm 0.01$ | $0.94 \pm 0.01$ | $0.62 \pm 0.01$ | $0.04 \pm 0$ |
| $\varphi_{TL}$ (w) [°] | $71 \pm 2.7$ | $79.9 \pm 3.3$ | $92.9 \pm 3.2$ | $103 \pm 4$ | $109.2 \pm 3.7$ |
| $\varphi_{TL}$ (t) [°] | $72.2 \pm 2.5$ | $83 \pm 2.8$ | $96.1 \pm 3.8$ | $106.5 \pm 4.3$ | $112.2 \pm 2.9$ |
| $M_{TL}$ (w) | $0.000 \pm 0.012$ | $-0.033 \pm 0.032$ | $-0.087 \pm 0.035$ | $-0.091 \pm 0.037$ | $-0.015 \pm 0.006$ |
| $M_{TL}$ (t) | $0.004 \pm 0.005$ | $-0.027 \pm 0.041$ | $-0.121 \pm 0.057$ | $-0.111 \pm 0.043$ | $-0.015 \pm 0.01$ |

Mean values ± SD for the axial force ( $F_{aTL}$ ), the leg length change ( $\Delta l_{TL}$ ), the angle between leg and trunk vector ( $\varphi_{TL}$ ), and the hip torque ( $M_{TL}$ ) at walk (w) and at trot (t). Note that $F_{aTL}$ , $\Delta l_{TL}$ and $M_{TL}$ are dimensionless. $l$ is leg length and $l_0$ leg length at TD, $m$ = mass and $g$ = gravity.

### Thoracic limb, tangential leg function

As for the pelvic limbs, the thoracic limbs of all breeds touched down with a mean angle of attack φ_0_ of about 70° at walk. Maximal mean retraction angles φ_PL_ were about 115° for Beagles and Malinois, 109° for Whippets and ∼107° for French Bulldogs (Tab. 3). The torque pattern displayed by all breeds was biphasic. However, the first retraction phase was rather short. It follows protractor torque until TO (Toe off). The work exerted about the scapular pivot was negative (see Fig. 2D).

At trot, the leg touched the ground at steeper angles than those observed for walk. However, differences are not significant (for gait, breed or gait*breed *p* > 0.05). Except the Beagle, the other three breeds retracted their thoracic limb during stance phase at trot more than at walk (Tab. 3). Mean torque profiles look similar among breeds (Figs. 2D and 3D). The first positive half sinus (protractor torque) observed at walk almost disappeared at trot. Therefore, all exerted tangential work was negative. Interestingly, Whippets showed a mean negative maximum torque, which was like the maximum torque exerted at walk. For the other three breeds, peak negative torque at trot was twice as large as at walk.

### Thoracic limb, VPP

The position of the thoracic limbs’ VPP (VPP_TL_) is gait and breed related, but the linear combination gait * breed effects was not significant. The VPP_TL_ was placed significantly (p < 0.001) higher at walk (for Whippets, Malinois and Beagles, about 0.7 while for French Bulldogs 1.15 of effective leg length at TD) compared to trot (all breeds showed different distances). Post hoc test revealed that only the VPP_TL_ obtained for the French Bulldogs was significantly different from the other breeds. The horizontal distance between VPP_TL_ and scapular pivot did not significantly vary between breeds nor did the linear combination gait effects * breed effects (see Tab. 4).

**Table 4.** VPP distances to the proximal pivots.

| Breed / Gait | dVPP<sub>PL</sub> [% l<sub>0</sub>] | dhVPP<sub>PL</sub> [% l<sub>0</sub>] | dVPP<sub>TL</sub> [% l<sub>0</sub>] | dhVPP<sub>TL</sub> [% l<sub>0</sub>] |
| --- | --- | --- | --- | --- |
| Beagle<sup>(1)</sup> / walk | 0.48 ± 0.17 | -0,19 ± 0,09 | 0.70 ± 0.08 | 0,15 ± 0,06 |
| Beagle<sup>(1)</sup> / trot | 0.24 ± 0.08 | -0,06 ± 0,03 | 0.54 ± 0.17 | 0,23 ± 0,03 |
| F. Bulldog<sup>(2)</sup> / walk | 0.92 ± 0.39 | -0,42 ± 0,15 | 1.15 ± 0.14 | 0,11 ± 0,12 |
| F. Bulldog<sup>(2)</sup> / trot | 0.26 ± 0.14 | -0,17 ± 0,07 | 0.92 ± 0.26 | 0,23 ± 0,09 |
| Malinois<sup>(3)</sup> / walk | 0.68 ± 0.16 | -0,13 ± 0,08 | 0.68 ± 0.14 | 0,14 ± 0,07 |
| Malinois<sup>(3)</sup> / trot | 0.18 ± 0.12 | -0,02 ± 0,06 | 0.25 ± 0.12 | 0,16 ± 0,04 |
| Whippet<sup>(4)</sup> / walk | 0.81 ± 0.21 | -0,09 ± 0,10 | 0.75 ± 0.19 | 0,26 ± 0,13 |
| Whippet<sup>(4)</sup> / trot | 0.18 ± 0.07 | 0,01 ± 0,04 | 0.34 ± 0.06 | 0,15 ± 0,08 |
| Gait | p<0.001(***) | p<0.001(**) | p<0.001(***) | n.s. |
| Breed | n.s. | p<0.01(**) | p<0.001(***) | n.s. |
| Gait*Breed | n.s. | n.s. | n.s. | n.s. |
| Post-Hoc-Tests: |  |  |  |  |
| 2\1 |  | P=0.018(*) | * |  |
| 3\1 |  | n.s. | n.s. |  |
| 4\1 |  | n.s. | n.s. |  |
| 3\2 |  | P<0.01(**) | * |  |
| 4\2 |  | P<0.01(**) | * |  |
| 4\3 |  | n.s. | n.s. |  |

dVPP<sub>TL</sub> (distance VPP- hip joint), dhVPP<sub>TL</sub> (horizontal distance VPP- hip joint), dVPP<sub>TL</sub> (distance VPP- scapular pivot) and dhVPP<sub>TL</sub> (horizontal distance VPP- scapular pivot) are dimensionless (% l<sub>0</sub>). Negative dhVPP<sub>PL</sub> values indicate that the VPP<sub>PL</sub> is located cranially to the hip joint. Positive dhVPP<sub>TL</sub> values indicate that the VPP<sub>TL</sub> is located caudally to the scapular pivot.

## Discussion

In the present work we analyzed dog global dynamics at walk and trot based on kinematic and single leg kinetic data recorded from French Bulldogs, Whippets, Malinois and Beagles. Because the four breeds analyzed differ in body proportions, mass, posture, and breed purpose, our results may inform about general principles of dog quadrupedal locomotion. We focused our work on the level of the effective leg. We analyzed both the axial and the tangential effective leg functions, and of whether those leg functions are related to each other via a local VPP controller.

We found that different to bipeds (Andrada et al., 2015a; Andrada et al., 2014; Maus et al., 2010) or facultative bipeds (Blickhan et al., 2018), and other than proposed by Maus and colleagues (Maus et al., 2010), dogs displayed two VPPs during walk and trot. The VPP of thoracic limbs (VPP_TL_) may serve as target of control and is located above and caudally of the scapular pivot, while the VPP of pelvic limbs, the VPP_PL_, is located above and cranially of the hip and may serve as control object of the pelvic limbs’ drive. Please consider, that the horizontal positions of both VPPs related to their proximal pivots largely influence both axial and tangential leg functions and leg work (Andrada et al., 2014; Blickhan et al., 2015). When the VPP is located above the proximal pivot as observed in human walking, the effective leg will likely display a rather symmetric kinematic behavior (similar inner leg angles at TD and TO), together with symmetric vertical GRF and protraction and retraction torques patterns. For such a configuration a simple spring-like axial leg behavior together with VPP-control is able to generate steady-state locomotion (Maus et al., 2010). If the VPP is located cranially to the proximal pivot, as known from birds or Japanese macaques and now depicted in dog pelvic limbs, the torque generated by the VPP control will generate only or mostly positive work. Birds and Japanese macaques use spring-damped effective leg functions to axially absorb energy and stabilize locomotion (Andrada et al., 2014; Andrada et al., 2015b; Blickhan et al., 2018). Interestingly, dogs, with the only exception of French Bulldogs during walk, displayed a different strategy. They extended their pelvic limbs during stance, indicating that energy was added to the system. Finally, if the VPP is placed caudally to the proximal pivot, as depicted in dogs’ thoracic limb, the torque generated will absorb energy. In this case the leg must add energy axially to the system for the sake of periodicity (Blickhan et al., 2015; Drama and Badri-Spröwitz, 2020). Accordingly, the dogs under study here, except Whippets during walk, add energy axially to the system. Taking this into account, it is not surprising that the axial leg function in dogs cannot be fully described with simple spring or spring-damper systems, although it was certainly more spring-like during trot. Our findings question several simple models based on spring-like or spring-dampened leg, e.g. (Herr and McMahon, 2001; McMahon, 1985; Nanua, 1992; Nanua and Waldron, 1995; Poulakakis et al., 2003; Poulakakis et al., 2006) and a well-accepted hypothesis like the strut limbs proposed by Carrier and colleagues (Carrier et al., 2005; Carrier et al., 2008).

Based on the principle that neuronal control is rather conservative, we hypothesized that locomotion control principles at the global level should be roughly similar among different dog breeds at the same gait. Our results could not falsify the above defined hypothesis and seem to support this idea. In general, gait changes influenced the position of the local VPPs and the patterns of the axial and tangential patterns more significantly than breed. However, as already mentioned, some breed related differences were found. In the following sections, we will discuss those differences, together with the influence of each VPP on limb control and dynamics.

### Pelvic limb control

The distance between pelvis and VPP_PL_ is gait and breed related. During walk and trot, the VPP is located cranially to the hip. Its distance to the hip was reduced in trotting dogs, permitting a more spring-like function of the effective leg. French Bulldogs exhibit significantly more cranially located VPP_PL_ during both walk and trot, which can be explained by their specific locomotion. This position induced only extensor torques in the hip and leg shortening during stance in contrast to the sinus pattern (extension-flexion) for the hip torque and leg lengthening observed for the other breeds in this study. Note that the latter patterns are also considered to be a general feature in dogs (Headrick et al., 2014; Ragetly et al., 2010) and in other quadrupeds during level locomotion, e.g. (Andrada et al., 2013; Witte et al., 2002). French Bulldogs display unusual pelvic limb three-dimensional kinematics as femoral abduction (>40°) and external rotation (>30°) during walk and trot (Fischer et al., 2018). This complex segmental kinematics may hamper leg lengthening, and therefore propulsion is only produced via hip torque. In addition, French Bulldogs have a more cranially located CoM, due to their relatively big head compared to their body. Accordingly, they exhibit lower pelvic vertical GRF/BW when compared to the other breeds analyzed in this study. This fact may permit them the use of more cranially located VPP_PL_ without significantly increasing hip torque, as shown in this work. A more cranially and higher located VPP_PL_ may permit faster and powerful movements also in non-sagittal directions e.g., during fighting (Carrier Fighting vs. running). In contrast, a more aligned and closer VPP_PL_ to the hip may display an optimum for striding locomotion. Accordingly, the Malinois and the Whippets display more aligned VPP_PL_ positions related to the hip. Finally, the cranial position of the VPP_PL_ explains the accelerating fore-aft GRF widely observed in quadruped locomotion. This is because accelerating forces are necessary to rotate the GRF to the more cranially located VPP_PL_ (see Fig. 5).

### Thoracic limb control

The distance to the VPP_PL_ from the scapular pivot (dVPP_PL_) is gait and breed related. Likely, this distance is reduced for trot as consequence of reduced joint torques and joint work as GRF increases. French Bulldogs exhibit significantly larger dVPP_PL_ for both walk and trot. Independent of gait, the VPP_TL_ is placed caudally to the scapular pivot. Interestingly, with the only exception of the Whippets, the horizontal distance between VPP_TL_ and scapular pivot is increased for trot, permitting larger energy absorption tangentially in the effective leg.

The more caudal position of the VPP_TL_ related to the scapular pivot explains the braking anterior-posterior GRF observed during most of the stance phase in quadrupeds (Andrada et al., 2013; Bertram et al., 2000; Budsberg et al., 1987; Fischer and Lilje, 2011; Lee et al., 1999; Riggs et al., 1993). While at TD and in the early support the leg placement adds to the braking A-P forces, during most of the stance, the protractor torque in the proximal pivot (necessary to rotate axial forces to the VPP_TL_) generates also negative A-P GRF. Thus, due to the more caudal position of the VPP_TL_, the thoracic limbs work against the retraction of the limb during the stance phase. This fact, which is counter intuitive, explains why the M. latissimus dorsi, the so-called main retractor of the humerus, actually remains silent during striding locomotion in dogs (Carrier et al., 2006; Carrier et al., 2008). Protracting torques in the scapula and/or in the shoulder joints computed via inverse dynamics were previously reported for dogs (Andrada et al., 2017), horses (Clayton et al., 1998; Colborne et al., 1998), small mammals (Witte et al., 2002) and rats (Andrada et al., 2013). However, the VPP control depicted in the present work explains the foundations of that torques and their relationships with the measured GRFs.

The large negative work produced tangentially is partially compensated by leg lengthening (axial positive work) during stance. This compensation is more marked during trot, in which Whippets, Malinois and Beagles axially compensate roughly 50%, and French Bulldogs more than 85% of the negative work absorbed tangentially. The question is why dogs (and perhaps all quadrupeds?) absorb energy tangentially and add energy axially in their thoracic limbs. The most satisfying response might be to deal with a more cranially located CoM. While the tangential energy absorption may prevent an uncontrolled thoracic leg retraction, the axial lengthening in the late stance may generate late skew vertical force profiles (vertical forces that are higher in the late stance, see Fig. 5). Early skew vertical force profiles for pelvic limbs and late ones for thoracic limbs are ubiquitous across quadrupeds, e.g., (Andrada et al., 2013; Fischer and Lilje, 2011; Manter, 1938; Merkens et al., 1986; Pontzer et al., 2014; Ren et al., 2010; Zumwalt et al., 2006). In previous works it has been pointed out that those vertical force profiles minimize the pitching moment around the center of mass and reduce limb-work (Jayes and Alexander, 1980; Usherwood and Granatosky, 2020). To reduce pitching moment, late vertical skew forces from the thoracic limb are low during early stance, because they act on a large moment arm around the CoM. However, to ensure weight support, the vertical force must increase through stance, as the moment arm about the CoM diminishes. The opposite is true for the pelvic limbs. Larger vertical GRF are exerted in the early support phase, as the moment arm is short, and they are diminished as the lever arm becomes longer (Usherwood and Granatosky, 2020).

### Outlook: the VPP concept provides a general control framework for legged locomotion

The relative motion between external markers and the underlying bone segment have been described as a major source of error in kinematic assessment (Günther et al., 2003; Reinschmidt et al., 1997; Torres et al., 2011). Those errors, however, were shown to be less significant when analyzing leg pro- and retraction as in our study (Fischer et al., 2018). Moreover, the fact that VPP control is still evident even when averaging data from all individuals of a breed (Fig. 4) indicates that the findings in this study are of general nature.

**Figure 4.**
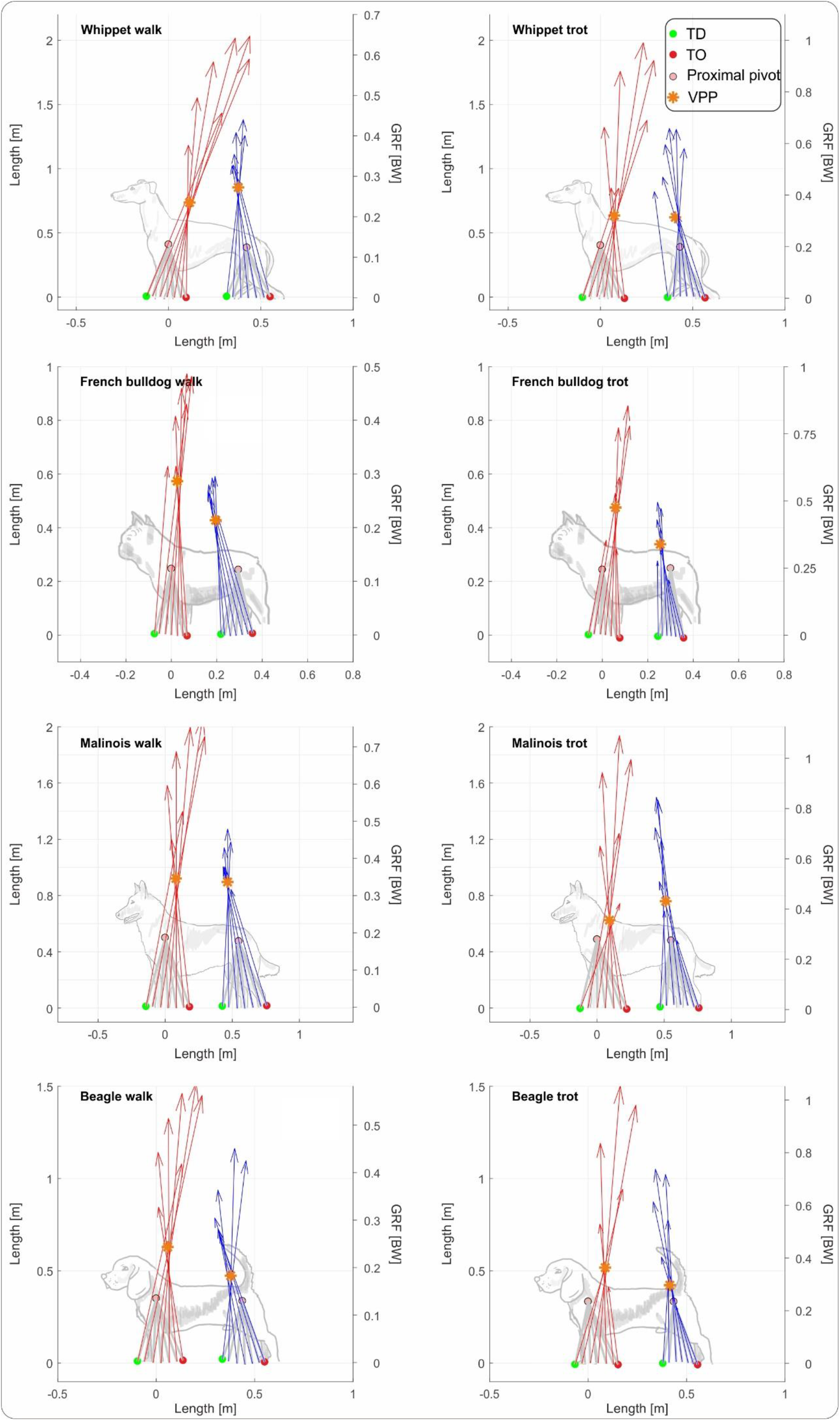
VPP control is still evident even when averaging data from all individuals of a breed. Subplots shows mean VPP_TL_ and VPP_PL_ for all analyzed breeds at walk and trot. Ground reaction forces and leg orientation are mean values for all individuals and strides of each bread. Red arrows correspond to the mean GRF of the thoracic limbs, while blue arrows correspond to the mean GRF of the pelvic limbs. Distal points of the legs and GRF were computed relative to the proximal pivots. Therefore, the proximal pivots can be displayed as fixed points (see methods). Superimposed dog sketches were included for an easier interpretation of the figures. They were not isometrically scaled. Beagle (N=5), strides walking = 84 for pelvic limbs and 77 for the thoracic limbs, strides trot = 104 for pelvic limbs and 85 for the thoracic limbs. French Bulldog (N=4), strides walking = 71 for pelvic limbs and 68 for the thoracic limbs, strides trot = 75 for pelvic limbs and for 49 the thoracic limbs. Malinois (N=4), strides walking = 35 for pelvic limbs and 42 for the thoracic limbs, strides trot = 34 for pelvic limbs and 28 for the thoracic limbs. Whippet (N=5), strides walking = 73 for pelvic limbs and 74 for the thoracic limbs, strides trot = 75 for pelvic limbs and 78 for the thoracic limbs.

**Figure 5.**
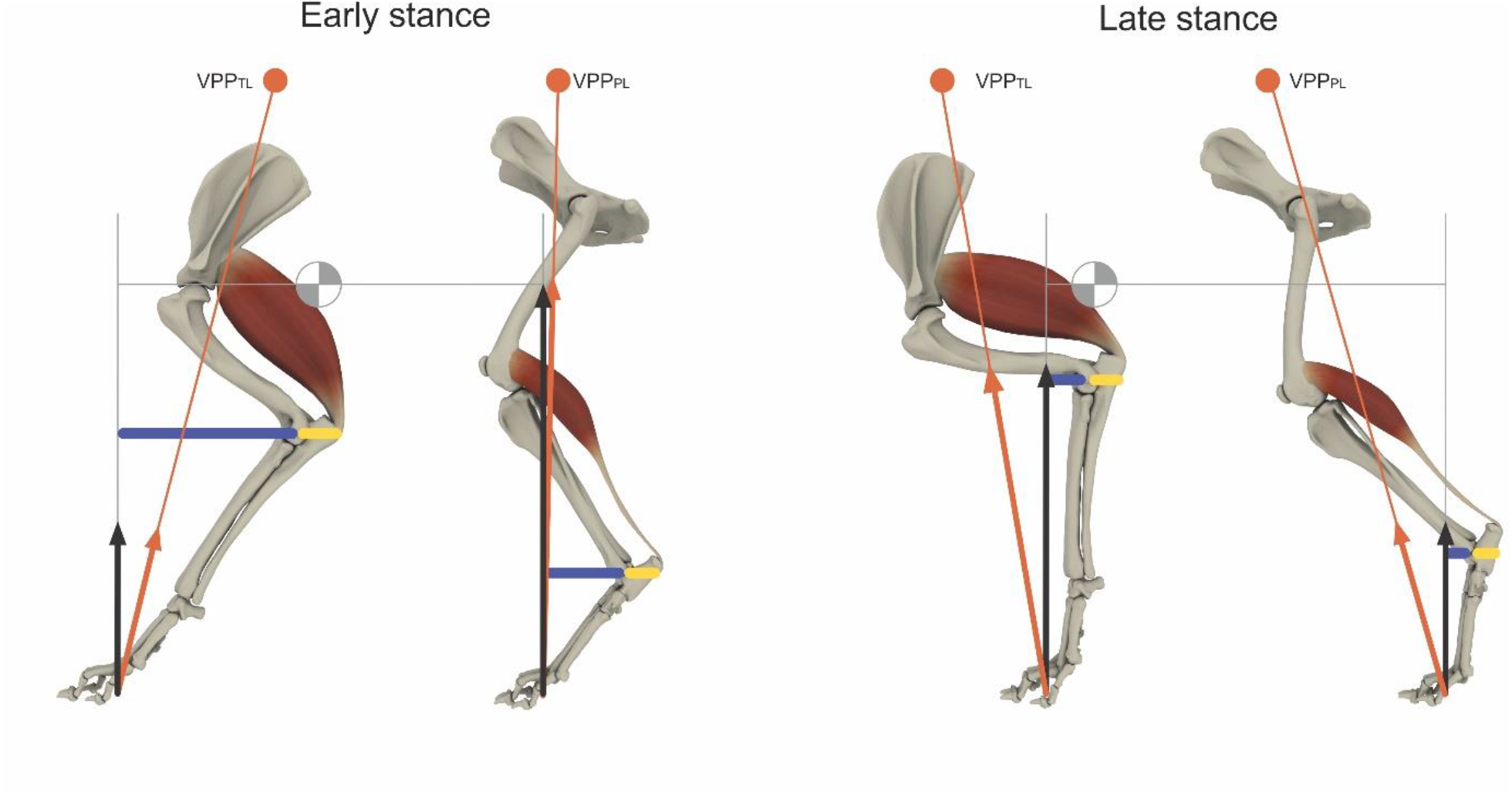
Skew vertical force profiles (black arrows) minimize the pitching moment around the center of mass and may reduce limb-work. Vertical forces from the thoracic limb are low during early stance, because they act on a large moment arm around the CoM. To ensure weight support, the vertical force must increase through stance, as the moment arm about the CoM diminishes. The opposite is true for the pelvic limbs. The caudal position of the VPP_TL_ relative to the scapular pivot permits tangential energy absorption during stance, while the rather cranial position of the VPP_PL_ relative to the hip generates energy tangentially. The key anti-gravity muscles, M. triceps caput longum and Mm. gastrocnemii, are preventing flexion by isometric contraction. Orange arrows represent the vector of the GRF.

From the present study, the VPP template emerges as the simple and general goal of global control for both bipeds and quadrupeds. The VPP template can perhaps be understood as a global representation of coupled contralateral central pattern generators (CPGs) (Brown, 1914; Grillner and Manira, 2020; Grillner and Zangger, 1979). Simulation studies showed that stable bipedal gaits emerge for certain values of VPP position in body frame and leg impedance (Andrada et al., 2014; Drama et al., 2020; Maus et al., 2010). Note that these parameters were adapted by our analyzed dogs when changing from walk to trot (see e.g., Tab. 4 and Fig. 4). We hypothesize that the tuning of VPP control parameters for a certain gait is learned during first months of life and possibly encoded in the coupling coefficients of the synaptic connections between CPGs. After this learning process, the coefficients may be set in a feedforward manner by excitatory and inhibitory command signals from higher control centers. The switch between quadrupedal and bipedal locomotion (from two to one VPP controller) might be performed by inhibitory commands from higher centers towards the most caudally located oscillators. Thus, under the VPP-framework, controlling both quadrupedal and bipedal gaits needs no new control parameters, but only tuning existing ones. This fact might explain why quadrupeds like Japanese macaques, bonobos, chimpanzees or sifakas perform facultative bipedalism without major problems.

## Acknowledgements

The authors thank Prof. Dr. Ingo Nolte for access to his motion laboratory for this experiment; Rommy Petersohn, Ben Witt (formerly known as Ben Derwel) and Lars Reinhard for technical assistance. Thanks to Christian Rode for always interesting discussions about the VPP. We thank the dog handling unit of Saxony’s police force (Sächsische Polizeihundestaffel) for providing the Malinois, Prof. Dr. Ingo Nolte for making available the Beagles, Dr. Holger Bunyan for helping us on several occasions with his Whippets, and Michael and Liane Lobback for coming with their pack of French Bulldogs.

## Availability of data and materials

The datasets used and/or analyzed during the current study are available from the corresponding author on reasonable request.

## Ethics approval and consent to participate

All experiments were approved by and carried out in strict accordance with the German Animal Welfare guidelines of the states of Thuringia (TLV) and Lower Saxony (LAVES) (Registration No. TLV Az. 22-2684-04-02-012/14, and 22-2684-04-02-009/15, LAVES 33.9-42502-04-14/1518, and 33.9-42502-04-15/1859).

## Authors’ contributions

E.A. conceived the study. E.A and M.S.F. conducted the experiments. E.A., G.H. analyzed experimental data, E.A., G.H., M.S.F. developed the figures. E.A., M.S.F. and H.W. grants acquisition. E.A. drafted the manuscript. All authors contributed to the interpretation of the results and revised the manuscript.

